# Automated and closed clinical-grade manufacturing protocol produces potent NK cells against neuroblastoma cells and AML blasts

**DOI:** 10.1101/2024.05.12.593780

**Authors:** Farhana Jahan, Leena Penna, Annu Luostarinen, Laurens Veltman, Heidi Hongisto, Kaarina Lähteenmäki, Sabine Müller, Seppo Ylä-Herttuala, Matti Korhonen, Kim Vettenranta, Anita Laitinen, Urpu Salmenniemi, Erja Kerkelä

**Affiliations:** Finnish Red Cross Blood Service, Research and Development, Helsinki, Finland; Finnish Red Cross Blood Service, Advanced Cell Therapy Centre, Vantaa, Finland; Finnish Red Cross Blood Service, Quality Management, Vantaa, Finland; Miltenyi Biotec B.V. & Co. KG, R&D Reagents, Bergisch Gladbach, Germany; Translational Cancer Medicine Research Program, University of Eastern Finland, Kuopio, Finland; University of Helsinki, Helsinki, Finland; Stem Cell Transplantation Unit, Helsinki University Hospital Comprehensive Cancer Center, Helsinki, Finland

**Keywords:** immunotherapy, Natural Killer cells, AML blast, CliniMACS Prodigy^®^

## Abstract

Natural killer (NK) cells have great potential as allogeneic immune cell therapy due to their natural ability to recognize and kill tumor cells, and due to their apparent safety. This study describes the development of an immunotherapy option tailored for high-risk acute myeloid leukemia (AML) in adults and neuroblastoma in children. A GMP-compliant manufacturing protocol for the local production of functionally potent NK cells is detailed in the study, including a comprehensive description of the quality control strategy and considerations for product batch specifications in early clinical development. The protocol is based on the closed, automated CliniMACS Prodigy® platform (Miltenyi Biotec) and a modified Natural Killer Cell Transduction (NKCT) process without transduction and expansion. NK cells are isolated from leukapheresis through CD3 depletion and CD56 enrichment, followed by a 12-hour activation with cytokines (500 IU/ml IL-2, 140 IU/ml IL-15).

Three CliniMACS Prodigy® NKCT processes were executed, demonstrating the feasibility and consistency of the modified NKCT process. A three-step process without expansion, however, compromised the NK cell yield. T cells were depleted effectively, indicating excellent safety of the product for allogeneic use. Phenotypic and functional characterization of the NK cells before and after cytokine activation revealed a notable increase in the expression of activation markers, particularly CD69, consistent with enhanced functionality. Intriguingly, even following a brief 12-hour activation period, the NK cells exhibited increased killing efficacy against CD33+ AML blasts isolated from patients and against SH-SY5Y neuroblastoma (NBL) target cells *in vitro*, suggesting a potential therapeutic benefit for AML and NBL patients.

## Introduction

Acute myeloid leukemia (AML) stands as the predominant form of acute leukemia in adults marked by significant clinical and genetic heterogeneity. While a subset of patients achieves a cure through a multidisciplinary approach combining chemotherapy, antibody-based treatment, and allogeneic hematopoietic stem cell transplantation (HSCT), the prognosis remains particularly poor for older individuals or those with relapsed or refractory AML (1). Neuroblastoma (NBL), on the other hand, is a challenging solid extracranial tumor in children. NBL is an extremely diverse disease, and for those with low or intermediate-risk NBL, the prognosis is generally favorable when treated with surgery and low-intensity chemotherapy (2). Despite high-risk NBL patients initially responding well to traditional therapy, a proportion of them ultimately succumb to recurrent disease (3). Therefore, there is a high demand to explore novel treatment options for both high-risk AML and NBL patients.

Natural killer (NK) cells are cytotoxic lymphocytes that hold great promise as an allogeneic immune cell therapy option. NK cell-based immunotherapy exploits their natural antitumoral activity (4). Importantly, as NK cells appear not to elicit graft-versus-host (GvH) reactivity, they are considered safe in allogeneic transplantation, even in the non-HLA-matched setting (5,6).

Many clinical trials in hematological malignancies using allogeneic NK cells have been published demonstrating a good safety profile and promising preliminary treatment responses. NK cells have been administered either alone (7–9) or in the HSCT setting (6,10–12). Cytokine activation has been included in the manufacturing protocol to improve the efficacy of NK cells. A landmark study by Miller *et al*. (13) already showed the feasibility and safety of CD3-depleted, overnight (O/N) IL-2 stimulated cells in AML. After that, NK cells or cell products with different stimulation protocols, mostly IL-2 (14–17) or a cytokine combination O/N (18,19) have been used in clinical trials, again along with HSCT or without it. Moreover, longer cultures with cytokines (20) or feeder cells have been used in the clinical setting (21–25). There are also a few clinical studies reporting on the treatment of NBL using haploidentical, enriched NK cells, demonstrating some therapeutic potential when combined with modern immunotherapy (dinutuximab/GM-CSF/IL-2) (26) or chemotherapy (27,28). The feasibility of treatment with IL-2-activated or expanded NK cells has also been demonstrated in NBL (29,30).

Taken together from published clinical trial papers, several hundred patients, both adult and pediatric, have been treated with NK cells in hematological malignancies, of which AML has been the most common diagnosis. Over 200 patients have received cytokine-activated NK cells. Moreover, over 100 NBL patients have received NK cells, either with or without an antibody targeted at GD2. These studies have demonstrated the convincing feasibility and safety of allogeneic NK cell therapy in AML and NBL.

In this manuscript, we describe the setup of a manufacturing protocol and quality control analytics for a locally produced, minimally manipulated, freshly administered NK cell product designed as a new immunotherapy option for Finnish cancer patients. We isolated and activated NK cells from healthy donor-derived leukapheresis products using an automated, closed, CliniMACS Prodigy® platform (Miltenyi Biotec) -based process, including CD3+ cell depletion, CD56+ cell enrichment, and cytokine stimulation (12-h) steps. Moreover, we isolated AML blasts from whole blood samples and assessed the efficacy of the activated NK cell products. The planned patient groups consist of adult high-risk R/R AML patients, and later on, pediatric poorly responding or relapsed NBL patients, i.e., cases where traditional treatment options are not available.

## Materials and methods

### Ethics statement

Donors and patients have given their written informed consents to participate and the study was conducted according to the principles of the Declaration of Helsinki. The starting materials for production were collected by leukapheresis at the Finnish Red Cross Blood Service (FRCBS, Helsinki, Finland) from healthy voluntary donors. The study protocol was reviewed and approved by the Regional Committee on Medical Research Ethics, Helsinki University Central Hospital (ethical approval number: HUCH/1492/2020). Batches of primary NK cells not produced in Prodigy® were obtained from surplus buffy coats from healthy, anonymous blood donors. Research permission was obtained from the local Blood Service Review Board (permission no.178-06/2023-01/2024, FRCBS, Finland). Whole blood (WB) samples from AML patients were collected at the Comprehensive Cancer Center, HUCH (ethical approval number: HUCH/12335/2022).

### Manufacturing and quality control workflow of NK cells in CliniMACS Prodigy®

The starting materials for production were collected by leukapheresis using the Spectra Optia® 61000 Apheresis System using a continuous mononuclear cell protocol (Version 11) (Terumo BCT). Donor eligibility was determined according to standard procedures defined in EU legislation for blood establishments. The leukapheresis products were stored at room temperature before further processing on the same day.

The workflow, including a sampling plan and a QC scheme are presented in Fig. 1. The manufacturing process, analytical tests, and tentative release criteria (S1 Table) for the different process steps and the end product lot specifications were designed based on the EMA guideline (31), the European Pharmacopoeia (Ph.Eur.), and on published literature. NK cell processing was performed at the Advanced Cell Therapy Centre grade D cleanrooms at the FRCBS.

**Fig 1.**
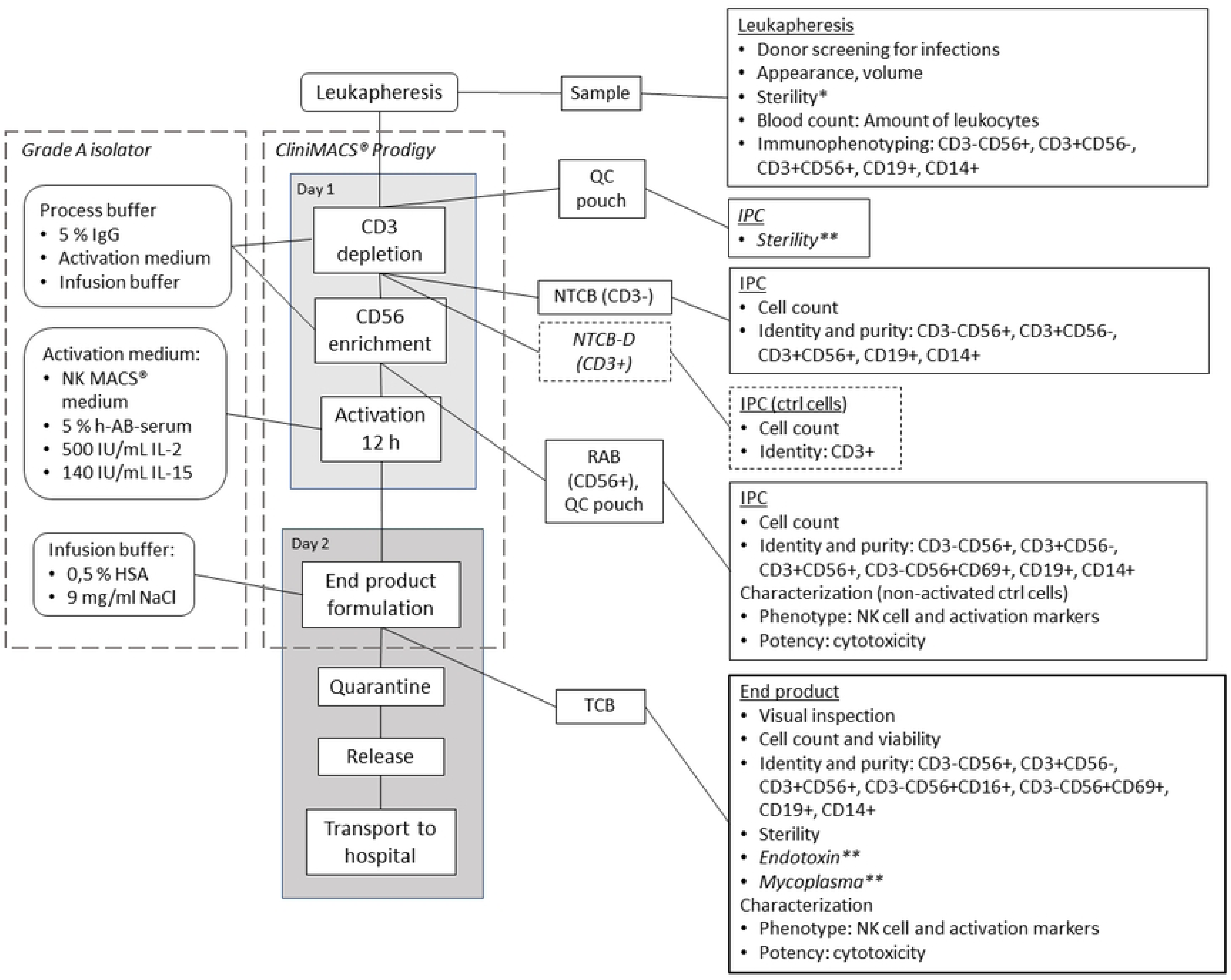
The workflow for the activated NK cell product in the CliniMACS Prodigy®. Manufacturing steps, sampling points and quality control (QC) analysis scheme are shown. Open phases are performed in the grade A isolator. *In the set-up phase, the sterility analysis was performed on the leukapheresis sample directly. **Analyses designed to be performed for the products proceeding to clinical studies but not performed in this study. IPC: in-process-control; NTCB-D: Non-Target Cell Bag Depletion; NTCB: Non-Target Cell Bag; NC: Nucleocounter; RAB: Re-Application Bag; TCB: Target Cell Bag

The open phases of the process, i.e., the preparation of media and buffers, were performed in a grade A isolator (Extract Technology Ltd) in grade D background according to GMP regulations. The isolator chamber was sterilized with vaporized H_2_O_2_, and particle monitoring was carried out during the whole operation time. For monitoring of aseptic conditions during the process, settle plates and glove prints were prepared on Tryptic Soy Agar (TSA) plates (Heipha) according to local standard operating procedures.

CE-marked reagents and consumables were used when available and all production materials were evaluated as suitable for manufacturing clinical products. The NK cell products were manufactured with the CliniMACS Prodigy**®** device using a modified, NKCT software program (at the time unpublished) and CliniMACS^®^ TS310 and TS520 tubing sets (Miltenyi Biotec). Depletion and enrichment were carried out using CliniMACS^®^ CD3 and CD56 reagents (Miltenyi Biotec), respectively, and published CliniMACS Prodigy® protocols. The NK cells were activated overnight (12-h) in the NK MACS^®^™ GMP Medium containing 5 % GMP human AB serum (Zentrum für Klinische Transfusionsmedizin Tübingen, Germany), 140 IU/mL MACS^®^™ GMP recombinant human IL-15 and 500 IU/mL MACS^®^™ GMP recombinant human IL-2. Activation was performed in a Prodigy CentriCult^TM^ chamber (part of the TS520 tubing set) under gentle shaking. The instrument setup for the activation utilized the NKCT protocol intended for CAR transduction and subsequent cultivation of NK cells, but only the cultivation, wash and harvest steps were used. The cells were harvested in saline containing 0.5% human serum albumin (Albunorm 200 g/l). The collected and analyzed cell fractions are outlined in Fig. 1. Aliquots of the NK cell end product were cryopreserved for later analyses.

### Sterility testing

After leukapheresis, 1 ml samples from the starting material, and at the end of the manufacturing process, 1 mL samples from the end products and 10 mL samples from the O/N culture media were analysed with BacT/ALERT VirtuO Microbial Detection System (BioMerieux) for aerobic and anaerobic bacterial growth.

### Cell count and viability

Cell samples were analyzed for the cell number and viability either with Nucleocounter NC-100 (Chemometec) or with trypan blue and TC-20 (Bio-Rad) automated cell counter. The complete blood cell counts of the leukapheresis starting materials were analyzed with Sysmex pocH 100i™ Hematology Analyzer (Sysmex Corporation).

From the GMP manufacturing process steps, samples were analyzed in duplicates and averaged. Log depletion was calculated as: log(# of T-cells/B-cells/monocytes in the start product / # of T-cells/B-cells/monocytes in the end product). The percentage cell recovery was calculated as: (total count of cells in target fraction / total count cells in original fraction) × 100%.

### Cell lines and primary NK cells

The cell lines used in this study, i.e., the K562-Luc2 cell line (ATCC® CCL243LUC2) and the SH-SY5Y NBL cell line (CRL-2266™), were obtained from the American Type Culture Collection (ATCC).

To confer luciferase expression, the SH-SY5Y cells were transduced with a lentiviral vector carrying the genes for enhanced green fluorescent protein (eGFP) and luciferase (Firefly luciferase, GenBank ID: M15077.1) separated by a P2A ribosomal skip sequence. The DNA was synthesized and cloned into the pLV-plasmid at Genewiz in Leipzig, Germany. Subsequently, 3^rd^ generation lentiviral vectors carrying the luciferase-eGFP gene were produced at the National Virus Vector Laboratory (the A.I. Virtanen Institute for Molecular Sciences, the University of Eastern Finland). The efficiency of transduction was assessed using flow cytometry.

The K562-Luc2 cell line was cultured in the IMDM medium, supplemented with 10% fetal bovine serum (FBS) and 8 µg/mL of blasticidin (all Gibco brand, from Thermo Fisher Scientific). The SH-SY5Y cell line was cultured in the minimum essential medium Eagle (EMEM, Merck) and F-12 nutrient mix (Gibco) medium, supplemented with 10% FBS, 2 mM L-glutamine, and 100 U/mL penicillin-streptomycin (Life Technologies).

Primary NK cells were isolated from peripheral blood mononuclear cells (PBMCs) in the laboratory (non-GMP) in order to add replicates for the cytotoxicity studies against AML blasts. First, PBMCs were separated from buffy coats using Ficoll-Paque Premium (GE Healthcare) density gradient separation. Subsequently, NK cells were isolated from PBMCs using Miltenyi Biotec’s NK Cell Isolation Kit following the manufacturer’s protocol. NK cells were cryopreserved for later use.

The isolated NK cells were maintained in the NK MACS Basal medium supplemented with NK MACS supplement (Miltenyi Biotec), 5% human AB serum (Sigma), 500 IU/mL of IL-2 (Proleukin S, 1 x 10^6^ IU/mL), and 140 IU/mL of IL-15 (Miltenyi Biotec). For experiments, NK cells were seeded at a density of 0.5 million cells per mL of media.

### Isolation and culture of AML blasts

The AML blast cells were isolated from whole blood (WB) utilizing MACSprep™ Chimerism CD33 MicroBeads (Miltenyi Biotec), following the manufacturer’s protocol. Concisely, 10 mL of blood was passed through a 30 μm nylon mesh to create a single-cell suspension. Subsequently, 500 μl of CD33 microbeads were added, and the mixture was incubated for 30 minutes at 4°C. Following incubation, CD33+ cells were isolated using the magnetic CD33 beads and eluted using an elution buffer obtaining the positively selected cell fraction.

CD33+ AML blast cells were cultured in the RPMI-1640 medium (Gibco), supplemented with a cytokine cocktail consisting of IL-3, IL-6, TPO, G-CSF, and SCF at a concentration of 20 ng/mL each (all cytokines from Miltenyi Biotec), along with 5% human AB serum (Sigma) (32–35). The culture was maintained by refreshing the culture media every 2-3 days. The cells were cryopreserved in a culture media containing 60% FBS and 10% DMSO (WAK Chemie medicals GmBH). The schematic diagram (Fig. 2) shows the overall procedure.

**Fig 2:**
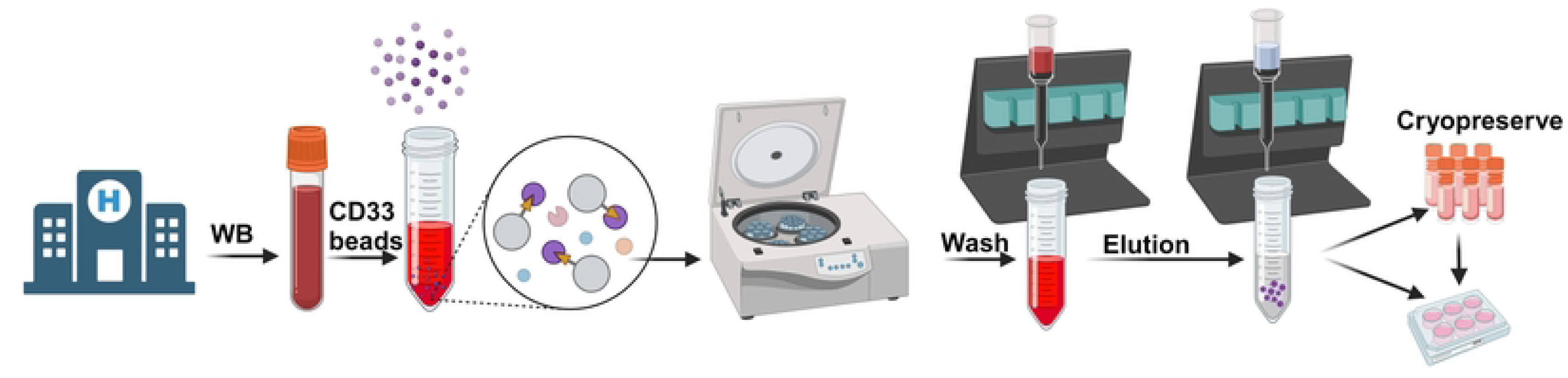
Flowchart illustration of the AML blast isolation process. The AML blasts are isolated from the whole blood (WB) of an AML patient for the *in vitro* experiments. Created with Biorender.com.

### Flow cytometry

#### QC panels

The NK cells from different manufacturing steps were characterized with flow cytometry (DxFLEX, Beckman Coulter Inc.) as indicated in Fig 1. using three antibody panels (S2 and S3 Tables). Antibody staining (all from Miltenyi Biotec) was performed according to the manufacturer’s instructions. Briefly, for starting material and in-process-control (IPC 1 and IPC 2) samples, 1×10^6^ cells were stained with the panel of antibodies in 100 µl of staining buffer containing 0.5 % human serum albumin (HSA, Albunorm, Octapharma) and 2 mM EDTA (ThermoFisher Scientific) in PBS (Invitrogen). For the starting material and the end product, duplicate samples were analyzed. For the end product, 2 x 10^6^ cells were stained and 1 x 10^6^ cells were collected for analysis. Moreover, CD3+ T cells were used as a positive control to set the gate when analyzing the NK cells from the end product. After runs 2 and 3, a T cell spike-in dilution series was performed to show the detection limit of the flow cytometry assay for residual T cells in the end product.

Cells were stained with isotype control antibodies to serve as a baseline for comparison in the analysis. For CD69 expression analysis, a fluorescence minus one (FMO) control was performed. To correct for spectral overlap between different fluorochromes the MACS® Comp Bead Kit for anti-REA antibodies was used. Additionally, 7-AAD staining (Miltenyi Biotec) was used to identify and exclude dead cells from the analysis.

#### Phenotype markers

The phenotype of NK cells was assessed immediately after the collection of the samples (before activation and after activation, i.e., after the CD56-enrichment phase and from the end product, respectively) with flow cytometry (S2 and S3 Tables). The staining protocol was essentially the same as for the QC panels, except that only 0.1 x 10^6^ cells were stained.

#### AML blasts

The purity of the AML blast cell fraction after isolation was examined by analyzing the expression of CD33 antigen on the surface of the blast cells. The flowthrough samples from the isolation columns (cells negative for CD33) were included for comparison. The studied surface markers are presented in the ‘AML blast phenotype’ panel (S2 Table). Blast cells (0.1-0.2 x 10^6^) were washed and blocked with Human TruStain Fc blocker (BioLegend) for 15 minutes at 4 °C. The samples were then stained with the fluorophore-conjugated antibodies or isotype controls in a flow cytometry buffer for 30 minutes at 4 °C in the dark and analyzed with the FACS symphony A1 (BD Biosciences).

#### Analysis

Flow cytometry results were analyzed with the FlowJo® version 10.0.7 or 10.8.0. software (FlowJo LLC, BD Biosciences). Results are shown as the percentage of positive cells from the parent population or as fluorescence intensity (for CD69). For the markers analyzed in duplicates, the averages were calculated. Examples of the gating strategies for leukapheresis starting material and activated NK cells (i.e., end product) QC analyses are shown in S1 and S2 Fig. Phenotype and activation markers were gated from the CD56-positive NK cells (gated from lymphocytes) based on their corresponding isotype controls.

### Cytotoxicity assay with cell lines and AML blasts

To measure the cytotoxic efficacy of the NK cells, the cells were co-cultured with K562-luc2^+^, or with luciferase-transduced SH-SY5Y NBL target cells, at several effector-to-target (E:T) ratios for 16-18 hours. The luciferin reagent (ONE-Glo luciferase reagent, Promega) was added and live target cells were quantified according to the manufacturer’s protocol with a Victor Nivo® multimode plate reader (Perkin Elmer). An equal number of the target cells alone was used to denote ‘hundred percent luminescence activity’.

Following the thawing of the AML blast target cells, a recovery period of 2-3 days was allowed in culture before proceeding with the cytotoxic assay. Both activated and non-activated NK cells produced in the lab and CliniMACS Prodigy® were employed as effectors. Effector cells were retrieved from liquid nitrogen storage. For non-activated conditions, NK cells were rested for 2 hours. For activated conditions, NK cells were incubated with cytokines overnight. The effector and target cells at various E:T ratios were co-cultured for 4 hours, after which the live cells were quantified using the CellTiter-Glo reagent (Promega). The cell titer reagent, measuring ATP, an indicator of cellular metabolic activity, was analyzed using a CLARIOstar Plus microplate reader (BMG Labtech), software version 5.20 R5. The resulting luminescent signal, directly proportional to ATP quantity, enables precise evaluation of cell viability.

### Degranulation assay

For the detection of target cell-induced degranulation of the NK cells, the cells were co-cultured with K562 or SH-SY5Y (NBL) target cells at a ratio of 2:1 for 4 hours in the presence of the degranulation marker lysosomal-associated membrane protein 1 (LAMP-1, CD107a) antibody (PE Vio-615 conjugated, clone REA792, Miltenyi Biotech) and after 1 hour of co-incubation, the GolgiStop™ protein transport inhibitor (BD Biosciences) was added. Surface expression of CD107a on NK cells was measured by flow cytometry. NK cells alone were employed as the reference for assessing the expression activation markers.

## Results

### Automated, three-step manufacturing process generated consistent yields of pure NK cells

We established a short, three-step production method for clinical production of activated NK cell product with the corresponding sampling scheme and quality control analyses for each process step (Fig. 1). The preliminary tests and specifications for the product designed for a clinical phase I/II trial are shown in Table S1.

Results of the flow cytometry analysis of the starting materials and corresponding CD3-depleted, CD56-enriched intermediate products as well as the end products are shown for three individual manufacturing processes P_NK1-3 (Fig. 3).

**Fig 3.**
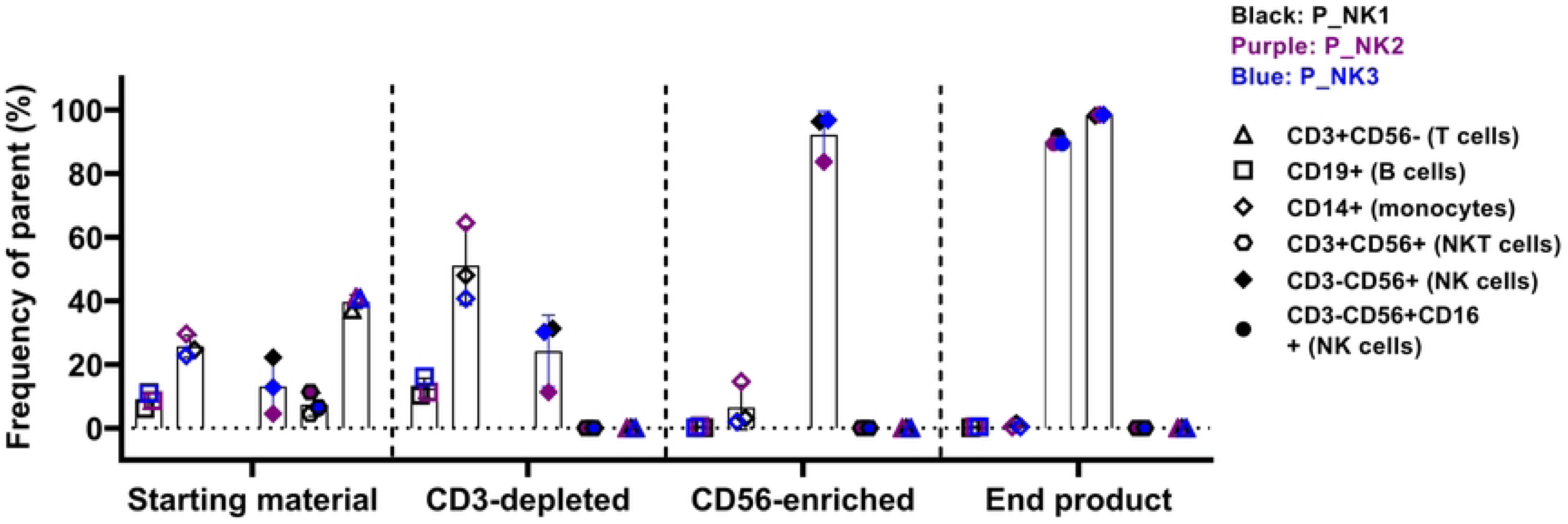
The distribution of cell populations at different process stages. The frequencies of different cell populations within CD45+ cells were assessed by flow cytometry at four distinct process stages: starting material, CD3-depleted, CD56-enriched, and end product. Different symbols represent the cell types: T cells (CD3+CD56-), monocytes (CD14+), B cells (CD19+), NK cells (CD3-CD56+) and two subsets of NK cells characterized by the presence or absence of CD16. The black, purple and blue color represent the three individual Prodigy^®^ processes (P_NK1-3). The bars represent means ±SD.

The number of different cell types and NK cell recovery percentage (yields) from the different process steps are shown in detail in Table 1. As a summary of three different manufacturing runs, the processes were very consistent yielding similar recovery percentages for NK cells (16.4%; 16.1%; 16.3 %, respectively), although the leukapheresed starting materials had very different cell compositions, representing donors having a high, low or medium frequency of NK cells (22.2%, 4.5%, 12.8%, respectively). The total NK cell numbers obtained from the processes, thus, varied (2.57 x 10^8^; 4.98 x 10^7^; 1.33 x 10^8^, respectively). The viability of the cells in the NK cell products were 95.4%, 88.6%, 93.5%, respectively.

**Table 1.** Cell numbers and NK cell recovery percentages from different process steps.

| Process 1 | WBC | NK cells |  |  | T cells |  |  |  | B cells |  |  |  | Monocytes |  |  |  |
| --- | --- | --- | --- | --- | --- | --- | --- | --- | --- | --- | --- | --- | --- | --- | --- | --- |
|  | total count | freq (%) | total count | recovery (%) | freq (%) | total count | recovery (%) | - logP | freq (%) | total count | recovery (%) | - logP | freq (%) | total count | recovery (%) | - logP |
| Leukapheresis | 7.08E+09 | 22.2 | 1.57E+09 |  | 37.3 | 2.64E+09 |  |  | 6.1 | 4.29E+08 |  |  | 24.6 | 1.74E+09 |  |  |
| CD3 depl. | 2.21E+09 | 31.4 | 6.93E+08 | 44.1 | 0.057 | 1.26E+06 | 0.05 | 3.32 | 10.3 | 2.27E+08 | 53.0 | 0.28 | 48.0 | 1.06E+09 | 60.8 | 0.22 |
| CD56 enrich. | 4.31E+08 | 96.3 | 4.19E+08 | 26.7 | 0.097 | 4.22E+05 | 0.02 | 3.80 | 0.16 | 6.96E+05 | 0.16 | 2.79 | 3.1 | 1.35E+07 | 0.78 | 2.11 |
| End product | 2.62E+08 | 98.2 | 2.57E+08 | 16.4 | 0.018 | 4.71E+04 | 0.00 | 4.75 | 0.185 | 4.84E+05 | 0.11 | 2.95 | 1.3 | 3.38E+06 | 0.19 | 2.71 |
| Process 2 | WBC | NK cells |  |  | T cells |  |  |  | B cells |  |  |  | Monocytes |  |  |  |
|  | total count | freq (%) | total count | recovery (%) | freq (%) | total count | recovery (%) | - logP | freq (%) | total count | recovery (%) | - logP | freq (%) | total count | recovery (%) | - logP |
| Leukapheresis | 6.88E+09 | 4.5 | 3.09E+08 |  | 41.1 | 2.83E+09 |  |  | 8.6 | 5.91E+08 |  |  | 29.6 | 2.04E+09 |  |  |
| CD3 depl. | 1.23E+09 | 11.3 | 1.39E+08 | 44.8 | 0.015 | 1.84E+05 | 0.01 | 4.19 | 11.3 | 1.39E+08 | 23.4 | 0.63 | 64.6 | 7.93E+08 | 38.9 | 0.41 |
| CD56 enrich. | 1.02E+08 | 83.8 | 8.53E+07 | 27.6 | 0.067 | 6.82E+04 | 0.00 | 4.62 | 0.61 | 6.21E+05 | 0.11 | 2.98 | 14.6 | 1.49E+07 | 0.73 | 2.14 |
| End product | 5.05E+07 | 98.6 | 4.98E+07 | 16.1 | 0.005 | 2.53E+03 | 0.00 | 6.05 | 0.45 | 2.27E+05 | 0.04 | 3.42 | 0.47 | 2.37E+05 | 0.01 | 3.93 |
| Process 3 | WBC | NK cells |  |  | T cells |  |  |  | B cells |  |  |  | Monocytes |  |  |  |
|  | total count | freq (%) | total count | recovery (%) | freq (%) | total count | recovery (%) | - logP | freq (%) | total count | recovery (%) | - logP | freq (%) | total count | recovery (%) | - logP |
| Leukapheresis | 6.37E+09 | 12.8 | 8.16E+08 |  | 40.9 | 2.61E+09 |  |  | 11.2 | 7.14E+08 |  |  | 22.9 | 1.46E+09 |  |  |
| CD3 depl. | 1.44E+09 | 30.2 | 4.35E+08 | 53.4 | 0.015 | 2.16E+05 | 0.01 | 4.08 | 16.1 | 2.32E+08 | 32.5 | 0.49 | 40.7 | 5.87E+08 | 40.2 | 0.40 |
| CD56 enrich. | 2.44E+08 | 96.8 | 2.37E+08 | 29.0 | 0.018 | 4.40E+04 | 0.00 | 4.77 | 0.20 | 4.89E+05 | 0.07 | 3.16 | 2.0 | 4.84E+06 | 0.33 | 2.48 |
| End product | 1.35E+08 | 98.5 | 1.33E+08 | 16.3 | 0.005 | 6.74E+03 | 0.00 | 5.59 | 0.45 | 6.06E+05 | 0.08 | 3.07 | 0.45 | 6.06E+05 | 0.04 | 3.38 |

311 *WBC: white blood cell*

The purity of the final NK cell product was high (CD3-CD56+ cells average 98.4%, SD 0.2) and the cells were mostly CD16-positive (average 90.3%; SD 1.6) (Fig. 3). T cells were depleted effectively (CD3+ cells average 0.01%; SD 0.007) (Fig. 3 and Table 1), as indicated also by log depletions (logP) of T cells (4.75; 6.05; 5.59, respectively) (Table 1). Thus, if dosing of NK cells would be 1 x 10^6^ cells /kg of the patient, the final T cell numbers would be clearly < 1000 cells /kg of the patient body weight in all the products. In addition to CD3-CD56+ NK cells, there were some monocytes (average 0.73 %; SD 0.48) and B cells left in the end products (average 0.36%; SD 0.15) (Table 1).

### Microbiological quality of NK cell production

No growth in either aerobic or anaerobic BactAlert cultivation was detected in any of the samples from the starting materials, final products or O/N culture media. No growth in any of the in-process control settle plates (9-12 plates/batch) from the isolator was detected, however, one single bacterial colony grew on one of the glove prints (8 plates/batch) from one process. The isolate was identified as *Rhodococcus* sp. The results from regular monthly environmental monitoring (active air samples and surface samples) of the isolator and the grade D background were all compliant with the requirements of ATMP-GMP. Likewise, non-viable particle monitoring and other monitoring of the clean room conditions revealed no other relevant non-conformances during the three NK cell production processes.

### Overnight activation protocol conserved a favorable NK phenotype

The NK cells were characterized extensively with flow cytometry for phenotype and activation markers both before and after the 12-h cytokine activation. The phenotype of the NK cells was not heavily affected by the 12-h cytokine activation based on the markers studied (Fig. 4), but donor-dependent changes in the expression levels of particularly CD57, NKG2A and NKG2D were observed. The expression of LAG3 and NKG2C was low in all samples.

**Fig 4.**
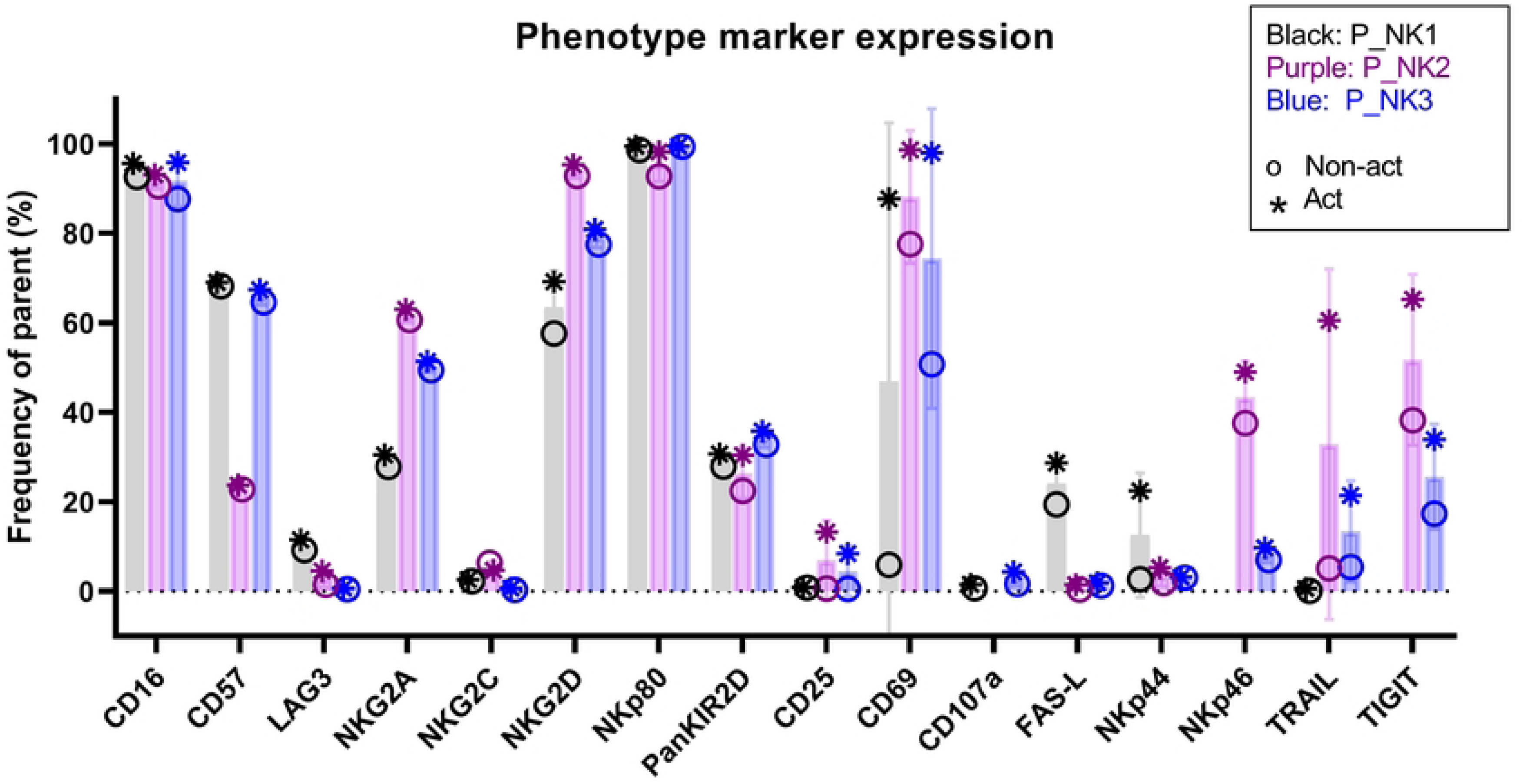
Phenotype marker expression in activated and non-activated NK cells. The expression of selected phenotype markers was assessed by flow cytometry from the CD56-positive lymphocytes. The black, purple and blue colors represent the three individual Prodigy^®^ processes (P_NK1-3) and the circle and star symbols stand for the non-activated and 12-h cytokine-activated NK cells. The bars represent means ± SD.

Early activation marker CD69 was upregulated in NK cells after the 12-h cytokine activation (Fig. 4). The expression of CD69 varied between donors before cytokine activation but showed upregulation in all processes after activation compared to the starting level. The expression of almost all activation markers was donor-dependent, as well as their reaction to cytokine activation. Donor NK2 had a higher expression of the activation markers in general, than donors NK1 or NK3.

Based on the activation marker expression of donor NK1, CD69 was chosen as a marker for successful cytokine activation and included in the QC panel from process 2 onwards. As CD69 was expressed also in the unactivated NK cells (from the CD56 enrichment phase), the increase in the fluorescence intensity was a more suitable indicator the activation of the NK cells (Fig. 5).

**Fig 5.**
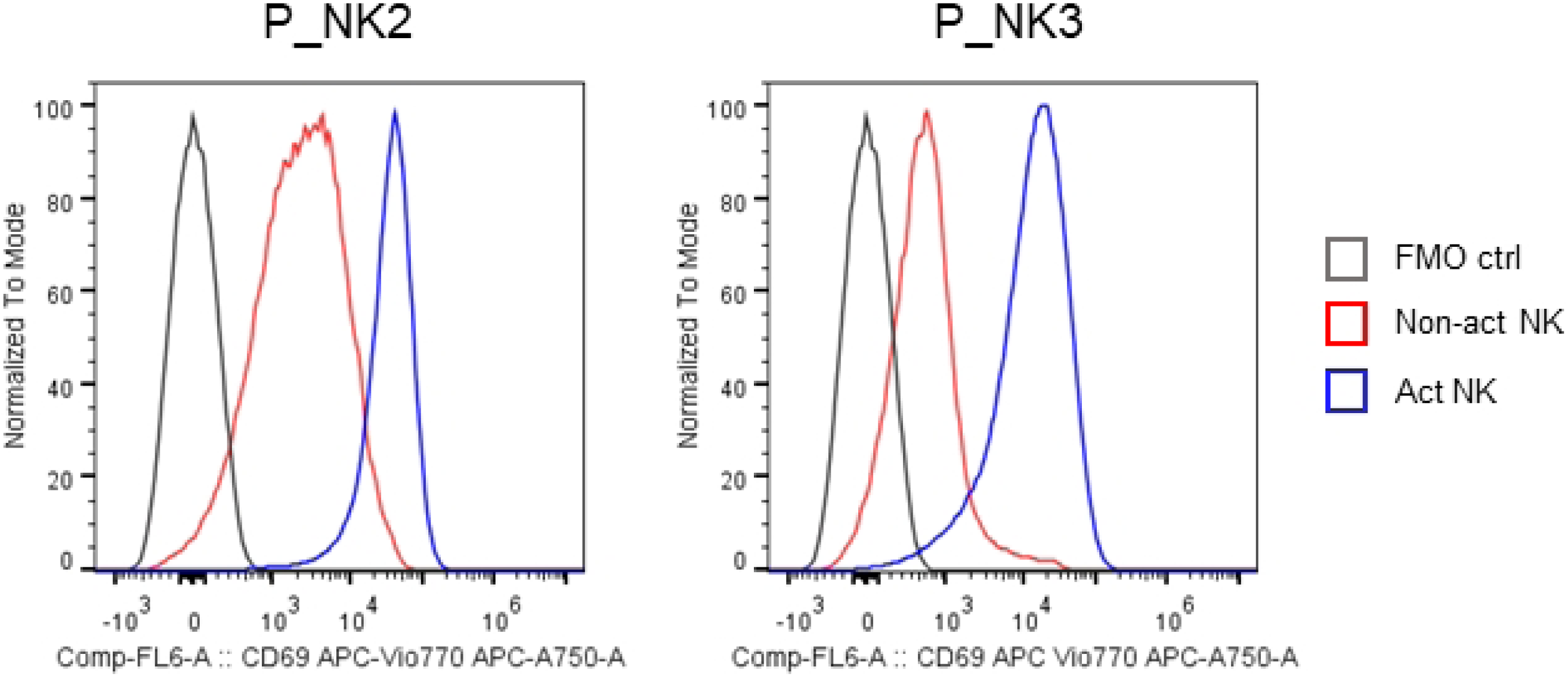
CD69 expression in non-activated and activated NK Cells. NK cells were produced in the CliniMACS Prodigy® and the expression was studied with flow cytometry from CD56+ NK cells, as part of the quality control panel. The fluorescence histograms for activated NK cells (end product), corresponding non-activated NK cells (after CD56-enrichment), and FMO control are shown.

### Successful isolation of AML blasts

Magnetic CD33 microbeads were chosen to facilitate the AML blast isolation. CD33 expression levels pre- and post-isolation were assessed with flow cytometry. The purity of CD33 was over 80% in both isolations (Fig. 6). Additionally, we analyzed the expression of other membrane molecular markers, including CD34, CD15, CD14, CD45, and CD45RA characteristic for AML blasts (Fig. 6). Intriguingly, differences in CD14 and CD45RA expression were observed between patients 1 and 2, potentially reflecting variation in the disease phenotype (36).

**Fig 6.**
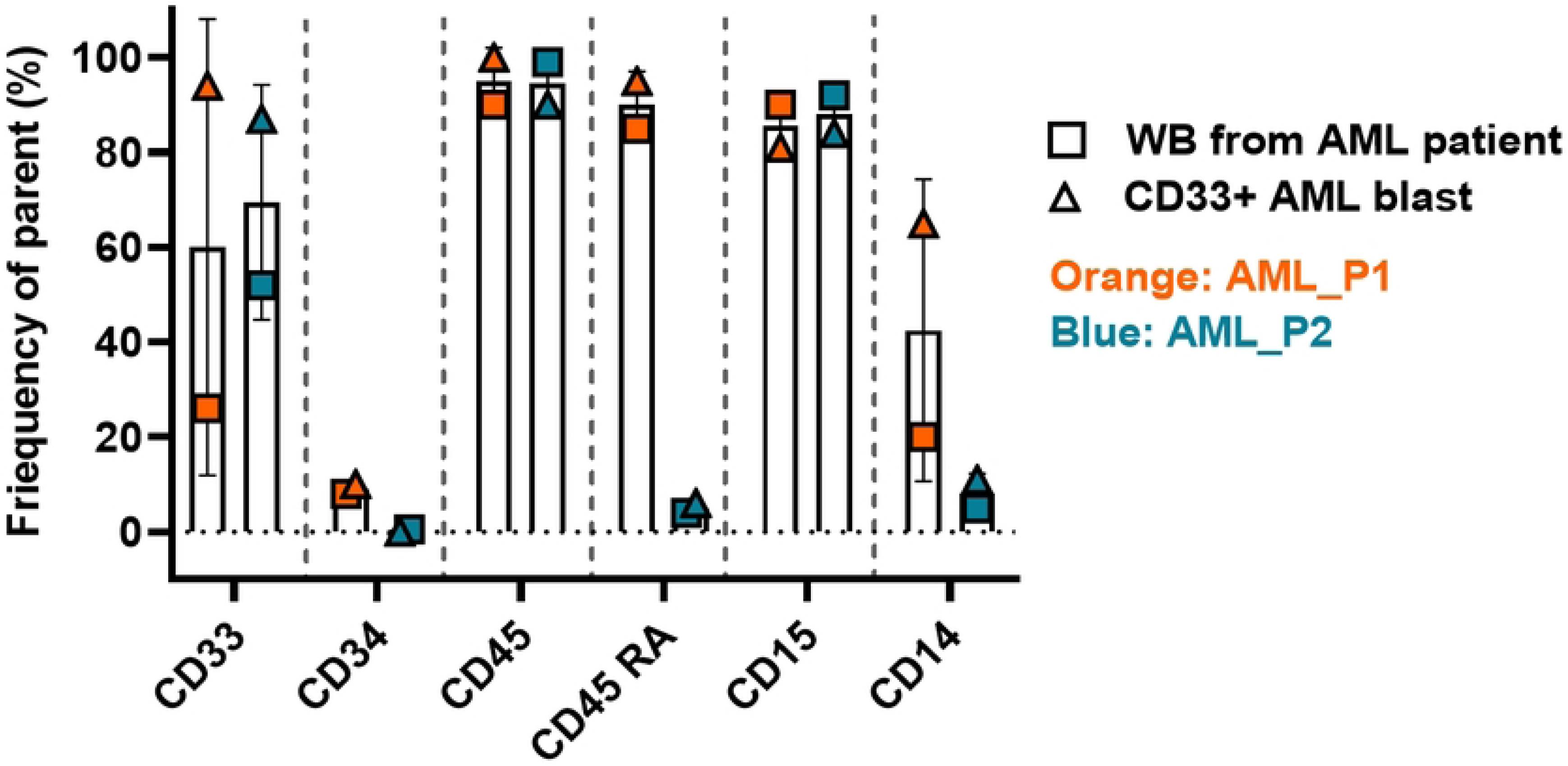
Expression of cell surface markers in AML blasts. The expression of cell surface markers was analysed with flow cytometry from cells collected from whole blood (WB) and after CD33 bead isolation from acute myeloid leukemia (AML) patient samples. The orange and blue colors represent the patients and the symbol expresses the sample type.

Patient selection for isolation and subsequent in vitro assays relied on a peripheral blood white blood cell count (WBC) threshold of ≤ 5×10^9^ cells per liter. This threshold was established after encountering difficulties in isolating CD33+ AML blasts, particularly in cases where the patient’s WBC count was less than 4 x 10^9^ cells per liter. Cell viability and growth were closely monitored every 2-3 days, although specific data are not presented here. Cryopreservation was performed with an expansion medium containing 60% fetal bovine serum (FBS) and 10% dimethyl sulfoxide (DMSO). This method yielded an AML blast viability of around 80%.

### NK cells showed high cytotoxicity against leukemia and neuroblastoma cell lines as well as primary AML blasts

The functional potential of NK cells from the manufacturing processes P_NK1-3 was analyzed using a luminescence-based cytotoxicity assay with the non-HLA-expressing universal K562 and the NBL-derived SH-SY5Y target cell lines. Intriguingly, even a short 12-h activation led to a higher NK cell killing efficacy against the SH-SY5Y and K562 target cells *in vitro*. Functionality was preserved even after a freeze/thaw cycle of the cells, although not on as high a level as that of fresh cells (Fig 7A,B).

**Fig 7.**
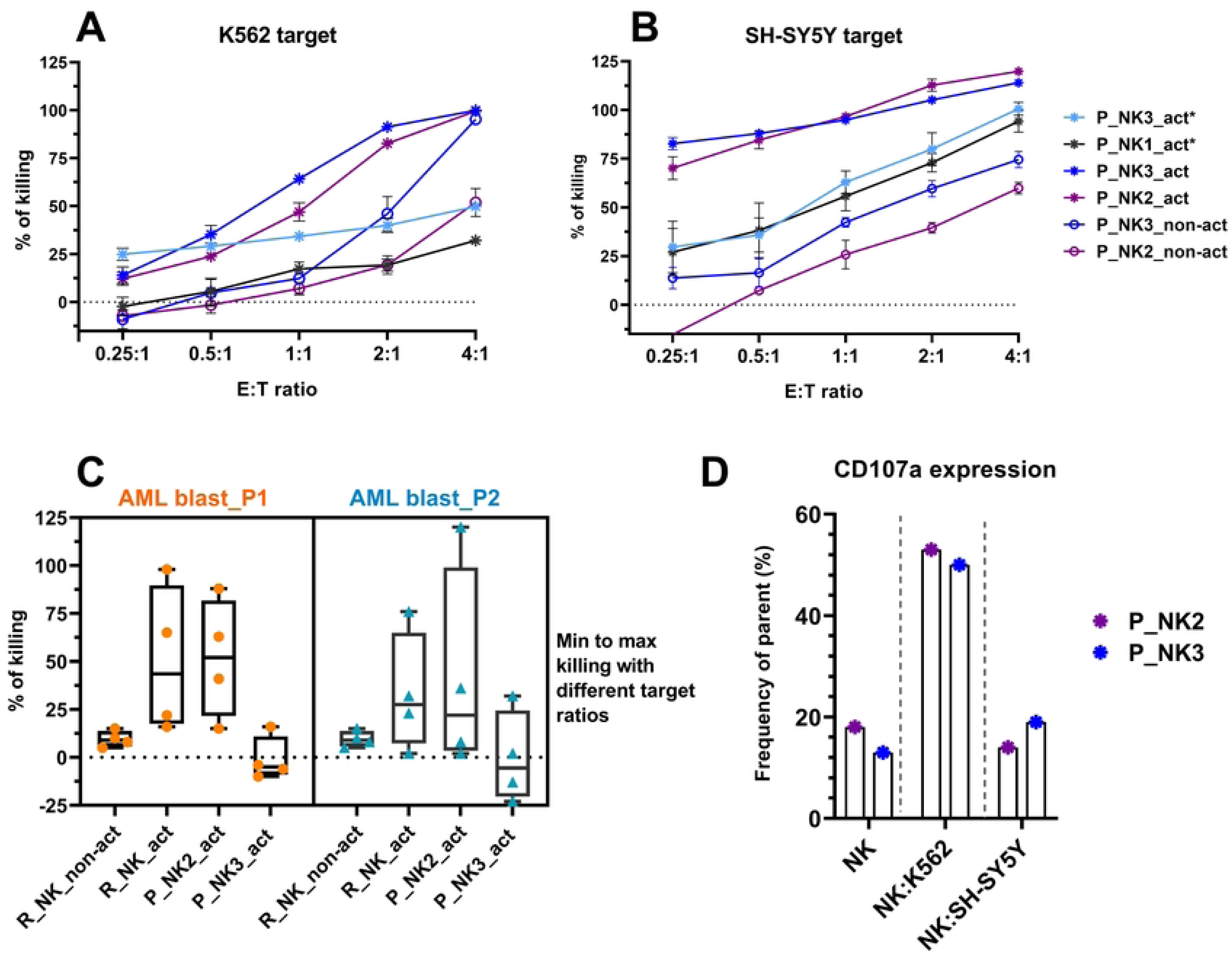
Functional assessment of NK cells post-production and activation. The luminescence-based assays compare the cytotoxic response of non-activated NK cells (‘non-act’) and 12-h cytokine activated NK cells (‘act’) against (A) K562 and (B) SH-SY5Y NBL targets at different effector to target ratios. Additionally, the P_NK3_act sample underwent post-activation characterization after a freeze/thaw cycle, denoted by an asterisk*. The P_NK1_act* sample was exclusively assessed post-freeze/thaw, with no corresponding non-activated control for comparison. The reported percentage of cell lysis represents the average of five replicates, with standard deviation (SD) indicated by error bars. C) The cytotoxicity of cytokine-activated NK cells targeting AML blast cells from patient samples. D) The degranulation response of NK cells following a 4-hour co-culture with K562 and SH-SY5Y cell lines, compared to NK cells cultured in the media alone (n=2). P_NK: NK cells were produced in the GMP CliniMACS Prodigy® conditions. R_NK and non-activated NK cells were produced in a non-GMP condition. AML blast_P denotes AML patient.

Next, the degranulation of the 12-h cytokine-activated NK cells was measured from co-cultures with K562 and with SH-SY5Y target cells. We observed a higher expression of CD107a in NK cells co-cultured with K562 cells for 4 hours than those co-cultured with neuroblastoma cells (Fig 7D).

Subsequently, the pivotal question was addressed of whether cytokine-activated NK cells possess the capability to target and eliminate blast cells derived from AML patients. To investigate this, AML blast cells were employed as the target, with non-activated NK cells obtained from anonymous healthy donors serving as the negative control in this experimental setup. Our findings revealed that the NK cells, whether activated in CliniMACS Prodigy® or in a non-GMP laboratory setting following a similar protocol, exhibited enhanced efficacy in eliminating AML blasts of two distinct patients when compared to non-activated NK cells (Fig 7C).

## Discussion

The objective of this study was to introduce a locally produced NK cell immunotherapy option tailored for Finnish cancer patients. Initially focusing on adult high-risk AML, we sought to extend this therapeutic approach to later include also pediatric neuroblastoma patients—addressing individuals with notably unfavorable prognoses. Utilizing the automated and well-established CliniMACS Prodigy® platform, we implemented a short cytokine stimulation (12-h) strategy for the NK cell activation, minimizing manual handling and mitigating potential sterility risks associated with open phases. Central to our approach was preserving the NK cells minimally manipulated prior to their potential administration to the patient, i.e., avoiding transduction procedures, freeze-thaw cycles, or long culturing periods.

The manufacturing process generating activated NK cells demonstrated substantial consistency and reproducibility across the three independent runs. Despite variations in the leukapheresis starting materials the processes consistently yielded similar recoveries of NK cells (16.1 - 16.4%). However, the three-step manufacturing protocol compromises the NK cell yield, which is in line with previous findings by others (16,37). The CliniMACS Prodigy® NKCT process is optimized for the transduction and 14-day expansion of NK cells (38,39), and not for a short period of cytokine activation only, which may also lower the cell yield. The optimal dose of NK cells remains unestablished and those in published clinical trials have ranged from 10^6^ to 10^8^ cells/kg of body weight. Some studies have shown a trend for a better outcome in patients receiving higher numbers of NK cells (8,16), which has recently nudged the consensus towards higher NK cell doses. In a large clinical study by Bachanova and coworkers (15), the total NK cell dose did not correlate with the clinical response, nor did it in a smaller study by Romee *et al*. (18). However, in clinical trials studying NK cells, different manufacturing methods and distinct treatment regimens have been applied, rendering dosing correlations complicated. It is important to note that the optimal cell dose most likely depends on various factors such as the type and stage of cancer, source and quality of NK cells, conditioning regimen, cytokine support, and combination with other treatments. The possibility for an optimal, alloreactive donor selection for more efficacious NK cells might mitigate the need for a high number of cells (40,41).

The purity of the NK cell product was consistently high with CD3-CD56+ cells averaging 98.4%. Effective depletion of T cells minimizes the potential for untoward immune responses (6). The purity of the final product was considered essential, as we were also aiming at treating pediatric neuroblastoma and wanting to eliminate T cell or B cell–related complications. Depletion of T cells was indeed very effective in all the production runs, average residual T cell percentage being 0.01%. With dosing of 1 x 10^6^ cells/kg patient weight the average residual T cell number would be 100 cells/kg, which based on published clinical trials can be considered very low.

The observed expression patterns of phenotype markers, including CD16, CD57, NKG2A, NKG2D, LAG3, and NKG2C, suggest a stable phenotype of the NK cells throughout the manufacturing process (42). Donor-dependent variation in the activation marker expression was evident, already before cytokine activation. There was also variation in the CD69 expression, nevertheless, CD69 was systematically upregulated already after 12-h cytokine activation. Thus, CD69 was included as a surrogate marker of successful activation in the QC panel of the final product, as due to time restraints, potency assay cannot be conducted for a freshly administered product before release for infusion. The measurement of NK activity through CD69 upregulation has been shown to correlate with NK cytotoxicity and degranulation (43,44).

Following a short 12-hour cytokine activation, the cytotoxicity assays conducted on the NK cells produced in the three CliniMACS Prodigy® runs revealed a notable enhancement in the killing efficacy against the NBL-derived SH-SY5Y and K562 target cells. The functionality of these activated NK cells was retained over a freeze/thaw cycle, albeit not at the same level as observed in fresh cells (Fig. 7A, B). IL-2 and IL-15 are known for their vital role in NK cell biology and are known to promote the maturation and survival of NK cells. Specifically, the *in vitro* exposure of NK cells to these cytokines, either alone or in various combinations, has been shown to enhance their cytotoxic activity against various target cells (37,45–50), which is in line with our results using a short cytokine activation.

The expression of CD107a in NK cells has been linked to NK cell degranulation and cytotoxic activity, particularly in response to IL-2 stimulation. The degranulation response is stable over time and does not affect the long-term viability or killing potential of NK cells (51). The higher expression of CD107a in NK cells during interactions with K562 cells compared to the neuroblastoma-derived SH-SY5Y cells in a 4-hour co-culture suggests a differential degranulation response based on the target cell type. As our NK cells readily killed the SH-SY5Y target cells at the later time point with the longer (16-18h) co-incubation, the lower level of degranulation at 4 hours may reflect diverse reaction kinetics in cell recognition and granule release, or imply NK cells relying on distinct killing mechanisms when encountering different target cell types (52).

A central aspect of the characterization methodology in this study was the isolation of AML blasts from the whole blood samples using CD33 microbeads, as well as subsequent cryopreservation and successful resuscitation of the isolated blasts. The use of CD33 microbeads for AML blast isolation is substantiated by the prevalent expression of CD33 in AML blast cells and its distinct nature as an AML antigen (53). The consistent CD33 purity levels obtained (over 80%) emphasize the efficacy of our isolation method (Fig. 6). This method provides an employable opportunity to investigate the killing efficacy of an NK cell product in a clinically relevant context using the actual donor-patient pairs from the clinical phase. CD33 bead selection from a whole blood sample offers convenience, decreases processing time and resource requirements as well as diminishes unwanted interference from other cell types, compared to previously used AML blast collection methods (18,54), e.g. blast cell aspiration from the bone marrow of AML patients, or sorting of blast cells from Ficoll-isolated PBMCs.

Investigation of cytokine-activated NK cells to target isolated AML blasts from patients demonstrated markedly enhanced efficacy compared to non-activated NK cells, supporting their therapeutic utility in AML treatment (Fig. 7C). Although statistical significance could not be established due to a limited number of replicates, the consistent trend across multiple runs supports the observed effect.

This study also served as an initial characterization of the aseptic process of NK cell production. The main part of the production process was completed in the CliniMACS Prodigy^®^ equipment, which is a closed system optimized for preserving aseptic conditions. The short open phases were performed in an isolator, which, according to process controls, completely retained class A requirements except in one process, where we detected a single *Rhodococcus* sp. colony in one glove print sample. *Rhodococcus* species are common environmental microbes, and as corrective and preventive actions, we improved our disinfection procedures during the uptake of production material into the isolator. Results from BactAlert cultivation showed that after each process, microbiological quality of the final product and the O/N culture media were according to specifications, as no growth of bacteria was detected. The automated growth-based BactAlert method (Ph Eur 2.6.27) is highly sensitive in microbial detection, but requires a 7 day-incubation period, and thus, release of the NK cell product would need to be done before obtaining the final results from sterility testing. It is therefore pivotal to optimize and control the aseptic process at all levels. As a next step, complete validation of clinical-grade production would require aseptic process simulation (APS), a routine procedure in ATMP validation. Readiness for clinical-grade production would also need the establishment of endotoxin and mycoplasma testing of the final product.

Limitations in the study include the low number of replicates. Only three product manufacturing runs in GMP conditions were conducted. The results from the quality control analyses were, however, consistent, including the composition, purity and cell yield. Moreover, to mitigate the lack of GMP NK cell samples, non-GMP-produced NK cells from buffy coats were additionally studied to address the effect of cytokine activation on the efficacy of NK cells against AML blasts. Our emphasis was on streamlined manufacturing process and purity. Due to low NK yield achieved with our protocol, modifications to the manufacturing are required to achieve sufficient cell numbers for adult patients. For future applications, one option to increase the cell yield could be NK cell expansion, doable in the closed CliniMACS Prodigy® system, with the production protocol we have used here. It is good to acknowledge, however, that there are indications that long expansion, either with cell lines or cytokines, might make NK cells quickly dysfunctional of exhausted *in vivo*, despite being highly active *in vitro* (55). To produce optimal NK cells for patients, further understanding on NK cell physiology and culture conditions is thus required.

## Conclusion

In this study, we established a GMP protocol for the local manufacturing of functionally efficient NK cells utilizing the CliniMACS Prodigy® platform. Moreover, a comprehensive description of the QC strategy, including the analytical tests, sampling points, and considerations for product batch specifications in early clinical development are described. Our detailed characterization, including the established CD33+ AML blast *in vitro* method, show that the NK cells obtained were activated and efficacious *in vitro*, and their phenotype remained substantially unaltered. Due to the fairly low total cell number in the final product, effective doses could only be achieved for pediatric patients and further protocol improvements would be needed.

## Acknowledgments

First, we are deeply grateful to the patients and leukapheresis and blood donors for their kind contribution to the study. Miltenyi Biotec is acknowledged for the technical support, and special thanks to Henrik Moller Thomsen for continuous support and training. We want to thank Advanced Cell Therapy Centre (ACTC) laboratory technicians for performing the CliniMACS Prodigy® process runs and clean room maintenance. The microbiology team is acknowledged for the microbiological sample analysis. Diana Schenkwein, Jan Koski, Solja Kalha and the laboratory personnel are acknowledged for their contribution to designing the lentiviral vectors and for the luciferase-positive target cell line generation.

## Author contributions

Conception or design of the work: FJ, MK, KV, ALu, US, EK; Acquisition, analysis, or interpretation of data: FJ, LP, ALu, LV, HH, KL, MK, ALa, US, EK; Provision of study material: SYh; Primary writing of the manuscript: FJ, EK; Editing of the manuscript: LP, ALu, HH, KL, SM, MK, KV, ALa; Approved the submitted version: FJ, LP, ALu, LV, HH, KL, SM, SYh, MK, KV, ALa, US, EK.

## Supporting information

**S1 Fig. Gating strategy for the leukapheresis starting material.** Example of the gating strategy applied for all leukapheresis starting materials. Similar gating strategy was applied for the samples after CD3 depletion.

**S2 Fig. Gating strategy for the end product.** Example of the gating strategy applied for all the activated NK cell products. Similar gating strategy was applied for the samples after CD56 enrichment, with the exception of CD16, which was not studied from the CD56 enriched samples.

**S1 Table. Tentative quality control tests and specifications for NK cell products planned for clinical phase I/II trial.**

**S2 Table. The list of all antibodies used in the study.**

**S3 Table. The flow cytometry panels and configurations for quality control analyses.**

